# Subset-specific and temporal control of effector and memory CD4+ T cell survival

**DOI:** 10.1101/2023.03.01.530323

**Authors:** Sharmila Shanmuganad, Autumn Ferguson, Aditi Paranjpe, Eileen Elfers Cianciolo, Jonathan D. Katz, Marco J Herold, David A. Hildeman

## Abstract

Following their proliferative expansion and differentiation into effector cells like Th1, Tfh, and T central memory precursors (Tcmp), most effector CD4+ T cells die, while some survive and become memory cells. Here, we explored how Bcl-2 family members controlled the survival of CD4+ T cells during distinct phases of mouse acute LCMV infection. During expansion, we found that Th1 cells dominated the response, downregulated expression of Bcl-2, and did not require Bcl-2 for survival. Instead, they relied on the anti-apoptotic protein, A1 for survival. Similarly, Th17 cells in an EAE model also depended on A1 for survival. However, after the peak of the response, CD4+ effector T cells required Bcl-2 to counteract Bim to aid their transition into memory. This Bcl-2 dependence persisted in established memory CD4+ T cells. Combined, these data show a temporal switch in Bcl-2 family-mediated survival of CD4+ T cells over the course of an immune response. This knowledge can help improve T cell survival to boost immunity and conversely, target pathogenic T cells.

## Introduction

Following their encounter with antigen in the context of self-MHC and appropriate co-stimulation, CD4+ T cells expand and differentiate into subsets of effector T cells with distinct function. Temporally distinct TCR, co-stimulation and cytokine signals not only drive the differentiation of these CD4+ effector subsets, but also modulate Bcl-2 family member expression and potentially, the survival of these subsets. However, *in vivo* mechanisms behind how activated CD4+ effector subsets survive during the expansion phase of an infection remain largely unexplored. After an acute infection is cleared, although most CD4+ T effectors die, some survive as memory cells. The BH3-only protein, Bim is required to kill most effector T cells during the contraction phase of the immune response (1-3).However, some CD4+ T cells counteract Bim-driven death and become memory T cells. The mechanisms that control this cell fate decision, between apoptosis and long-term survival, remain unknown.

In contrast to CD4+ T cells, the heterogeneity of effector CD8+ T cells consists of short-lived effector cells (SLEC) marked by high expression of KLRG1 and low expression of CD127 and memory precursor effector cells (MPEC) which express low levels of KLRG1 and high levels of CD127. Further, we and others showed that most SLEC expressed low levels of Bcl-2, whilst most MPEC expressed higher levels of Bcl-2(4, 5). Importantly, a common gamma chain-STAT5-Bcl-2 network is required to combat Bim and limit the pool of effector T cells available to enter the memory compartment (6). In fact, the only cells capable of surviving in the absence of Bcl-2 are CD8+ T cells expressing very low levels of Bim (5).Thus, in effector CD8+ T cells, Bcl-2 is essential to antagonize Bim and promote the survival of MPEC into the memory pool.

Ours and others work showed that the anti-apoptotic molecule Mcl-1 is also critical for *in vivo* survival of antigen-specific CD4+ and CD8+ T cells (7, 8). The role for Mcl-1 in T cell survival is likely independent of Bim as the additional loss of Bax and Bak, but not the additional loss of Bim, rescues Mcl-1-deficient T cells *in vivo* (7). Although *in vitro* studies have suggested a role for Bcl-xL in CD4+ T cell survival, *in vivo* experiments show that Bcl-xL is not required for blocking apoptosis of CD4+ T cells (9). Thus, the anti-apoptotic Bcl-2 family members that contribute to CD4+ T cell survival during an immune response remain unclear.

Following acute bacterial and viral infection, CD4+ T cells differentiate largely into Th1, Tfh effector and T central memory precursors (Tcmp) (10-16). Interestingly, *in vitro* stimulation of CD4+ T cells leads to repression of Bcl-2 expression and upregulation of another anti-apoptotic Bcl-2 family member, A1(17). Unlike Bcl-2, A1 has three functional paralogues in mice (A1a, A1b and A1d)(18), and although the global loss of all three genes did not disturb CD4+ T cell homeostasis after viral infection(19), the antigen-specific effector CD4+ T cell response was not examined.

Here we used genetic and pharmacologic approaches to examine the role of Bcl-2 and A1 in the survival of effector CD4+ T cell subsets during distinct phases of acute lymphocytic choriomeningitis virus (LCMV) infection. During their proliferative expansion and differentiation, Th1 cells depended on A1 and not Bcl-2, whereas other subsets depended on both proteins for their survival. Additionally, antigen-specific cells in an EAE model also depended on A1 for their survival and showed reduced disease progression in the absence of A1, suggesting that A1 may play a critical role in effector Th17 survival. In contrast, during the contraction phase of the LCMV response, both Th1 and Tcmp cells relied on Bcl-2 to transition into the memory pool with Tcmp cells contracting markedly less than Th1 cells. Combined, our data demonstrate a temporal, subset-specific involvement of Bcl-2 family members to control CD4+ T cell survival at distinct phases of the immune response.

## Results

### Differential expression of Bcl-2 family members within CD4+ effector subsets

In response to antigen presented by professional antigen presenting cells, CD4+ T cells differentiate into cytokine secreting effector cells and undergo substantial clonal expansion (20-23). While most expanded CD4+ T cells die, some survive and become memory cells (24, 25). Mechanisms controlling the survival of effector CD4+ T cells remain unclear. To have an unbiased view of the expression of survival factors within various effector CD4+ T cell subsets, we infected wild-type(WT) mice with the Armstrong strain of LCMV, sorted splenic viral-specific CD4+ T cells at day 7 post infection(p.i.) using GP66:I-A^b^ MHC tetramers and subjected them to single cell RNA sequencing (sc-RNAseq) (26) (Fig 1A).

**Fig 1.**
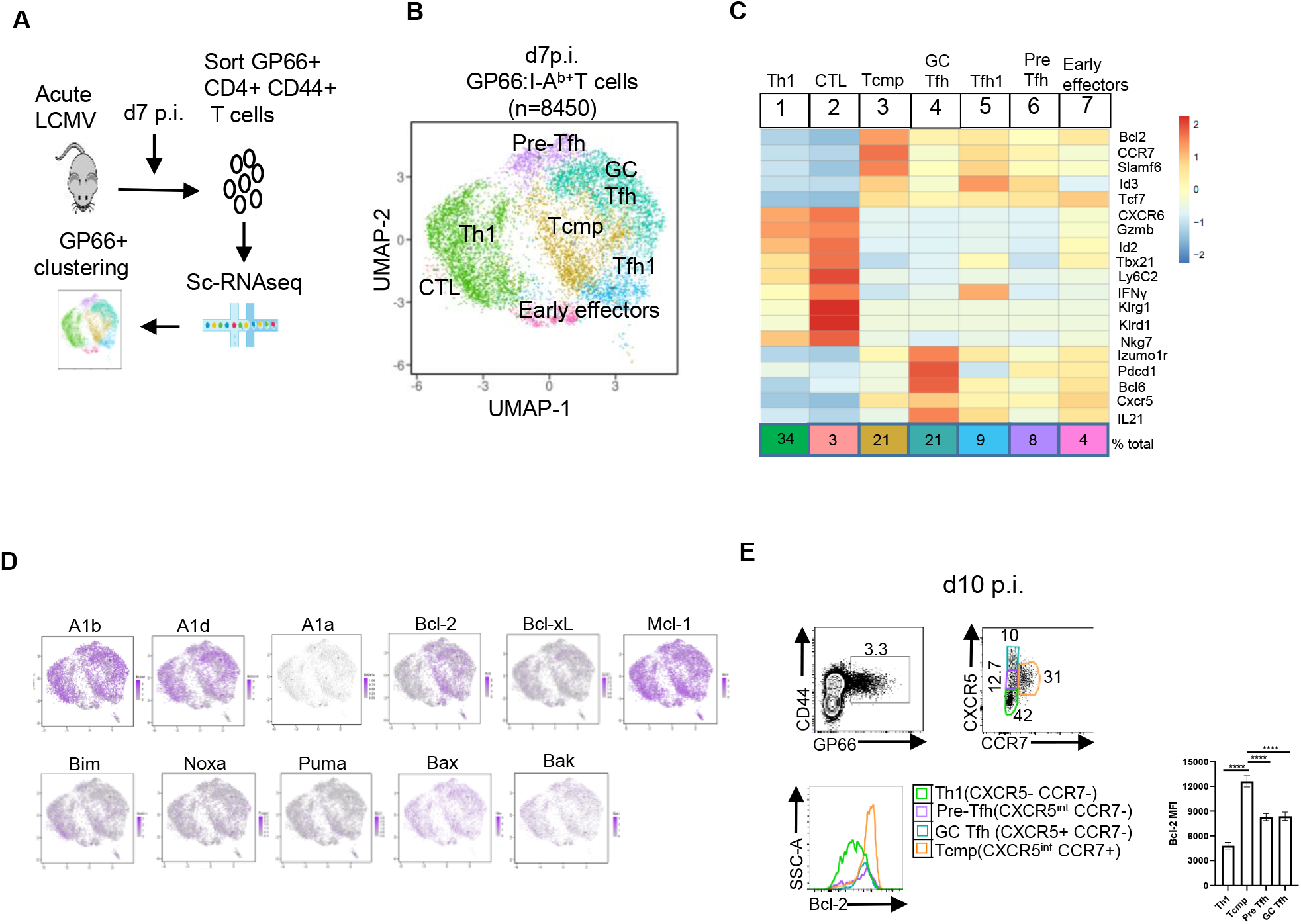
Expression of Bcl-2 family members in effector CD4+ T cells. (A) Outline of experimental procedure (B) UMAP plot of GP66-specific splenic CD4+ T cells from C57BL/6 (WT) mice at d7 post infection(p.i.) with LCMV showing clusters identified by variable gene expression. Each dot is an individual CD4+ T cell and colors highlight clusters. (C) Heatmap of Z-scores for the average expression of CD4+ T cell subset–specific genes in each cluster (D) Expression of Bcl-2 family members in clusters (E) GP66-specific CD4+ splenic T cells from WT mice at d10 p.i. with LCMV defined as Th1, Tcmp, GC Tfh and pre-Tfh subsets by the expression of CXCR5 and CCR7(top). Histogram and bar graph show Bcl-2 expression in these subsets(bottom). Data are representative of at least 3 independent experiments with n=3 or more mice per group and show mean ± SD. **** *p* < 0.0001 Student’s t test.

Dimensionality reduction analysis and unsupervised clustering revealed seven distinct clusters by Uniform Manifold Approximation and Projection (*UMAP*) visualization after regressing out cell cycle gene expression (Fig 1B). CD4+ T cell clusters were defined by lineage specific markers (Fig 1C) matching previously published data sets (14, 27). The clusters were largely divided into Th1, Tfh and T central memory precursors (Tcmp) subsets in agreement with the known heterogeneity of this population (12, 13). Cells in cluster 1 made up ∼ 35% of cells and readily identified as Th1 cells as they expressed known Th1 genes like CXCR6 and *Tbx21* (encodes TBET) (Fig 1C). Cells in cluster 2, comprised only∼ 3.5% of cells, shared expression of several Th1 genes, but were also enriched for cytotoxic markers like *Gzmb, Nkg7, Klrd1* and *Krlg1*(Fig 1C), so we designated this subset as CD4+ CTL (28). We identified cells in cluster 3 as T central memory precursors (Tcmp) as they had higher expression of *Ccr7, Slamf6, Bcl2* and *Id3* and accounted for ∼21 % of cells (Fig 1C) (10, 12, 14, 29). Cells in clusters 4 and 6 had higher expression of Tfh specific genes *Cxcr5, Bcl6, Icos, Pdcd1* (encodes PD-1; Fig. 1C) and together comprised ∼29% of cells. Of the Tfh clusters, cluster 6 had higher levels of *Selplg* (encodes PSGL-1) (Supplemental Fig 1A), and lower *Cxcr5* and *Pdcd1* expression compared to cluster 4(Fig 1C), hence we designated cluster 4 as Pre-Tfh (27) and cluster 6 as germinal center Tfh (GC Tfh) cells. Cells in cluster 5 comprised roughly 9.6% of cells and had gene expression similar to both Tcmp and Tfh, but also highly expressed *Tbx21* (encodes TBET) and *Ifng* (Fig 1C) and matched a previously described Tfh1 cluster (27). Lastly, cluster 7(∼4.1%) seemed to comprise early effector cells that had yet to make a fate decision based on their low expression of genes associated with several Th subsets (Fig 1C). We did not detect cells enriched for Treg-specific genes consistent with the lack of Tregs during LCMV acute infection at this time point (30).

Next, we assessed subset-specific expression of the major pro- and anti-apoptotic Bcl-2 family members. Interestingly, Bcl-2 showed a differential expression pattern with the highest expression in Tcmp, followed by Tfh and the lowest expression in Th1 cells (Fig 1D, Supplemental Fig 1B). Of three functional A1 genes (A1a, A1b and A1d) (18, 31), A1b and A1d were highly expressed across all clusters (Fig 1D, Supplemental Fig 1B). Bcl2l1 (encodes Bcl-xL) was high in Th1 and early effectors (Fig 1D, Supplemental Fig 1B). The expression of pro-apoptotic genes also varied among the clusters, with Bim being highly expressed in Th1 and Tcmp, but only modestly in all Tfh clusters. Puma, Noxa, Bax and Bak were expressed at varying levels across the different clusters, hinting at subset-specific roles for Bcl-2 family members in controlling apoptosis during expansion (Fig 1D, Supplemental Fig 1B). Further, flow cytometry analysis at d10 p.i. with LCMV of WT splenic GP66-specific CD4+ T cells using CXCR5 and CCR7 as markers to separate the different subsets, revealed three main subsets, consistent with prior work (10, 14) (Fig 1E). Tfh cells are split into GC-Tfh as they are CXCR5^+^ CCR7^-^ (∼10%) and also highly express PD1 and another population (CXCR5^int^ CCR7^-^, comprising ∼13%) that has been previously referred to as pre-Tfh cells (32) (Fig 1E). CXCR5^int^ CCR7^+^ cells (∼ 31%) will be referred to as T central memory precursors (Tcmp). CXCR5^-^ CCR7^-^ cells (∼ 40%), will be referred to as Th1 cells. Using these markers to closely approximate the sub-populations defined by the sc-RNAseq clusters, we analyzed their Bcl-2 levels by intracellular staining and flow cytometry. Strikingly, Th1 cells expressed significantly less Bcl-2 in comparison to Tcmp and Tfh clusters, recapitulating the sc-RNAseq data (Fig 1E). Thus, Bcl-2 family member expression differs substantially amongst effector CD4+ T cell subsets at the peak of the response.

### Subset-specific roles for Bcl-2 and A1 in CD4+ effector survival during expansion

We previously showed that Mcl-1 was essential for effector CD4+ T cell survival, however, it appears to largely independent of Bim (33). Further, Bcl-xL is not required for effector CD4+ T cell survival (9). Given the differential expression of Bcl-2 within effector subsets and the broad expression of A1b and A1d, we next examined the requirement of these proteins in the survival of effector CD4+ T cells. To avoid potential compensatory changes imposed by the requirement of Bcl-2 in naïve T cell survival (34), we bred Bcl2^fl/fl^ mice to CD4ER^T2^Cre transgenic mice (35) which express a tamoxifen-inducible Cre specifically within CD4+ T cells (referred to as Bcl2KO). To examine the role of A1, we crossed CD4ER^T2^Cre mice to recently developed A1a^-/-^b^fl/fl^d^-/-^ mice (36). In CD4ER^T2^Cre^-^A1a^-/-^b^fl/fl^d^-/-^ (referred to as A1DKO) mice, A1a and A1d are deleted in all cell types. In, CD4ER^T2^Cre^+^A1a^-/-^b^fl/fl^d^-/-^ (referred to as A1TKO) mice, A1b is deleted specifically in CD4+ T cells upon tamoxifen administration whereas A1a and A1d are deleted in all cell types.

Groups of wild type (WT), Bcl2KO, A1DKO, and A1TKO mice were infected with LCMV and tamoxifen was administered from d1 to d4 post infection to induce deletion of A1 and Bcl-2 (Supplemental Fig 2A) and splenic viral-specific CD4+ T cells were assessed 10 days p.i. using GP66:I-A^b^ tetramers(Fig 2A). First, the total numbers of GP66-specific CD4+ T cells was not changed in the absence of Bcl-2 (Fig 2B). On the other hand, the loss of either A1a, d or A1a, b,d resulted in significantly decreased numbers of GP66-specific CD4+ T cells (Fig 2B). The loss of GPP66-specific CD4+ effector T cells was not due to an impact of the global loss of A1a and A1d on GP66-specific naïve precursors (Supplemental fig 2B). Further, the frequency of CD44hi CD4+ T cells, which likely contain LCMV-specific T cells was significantly reduced in A1DKO and A1TKO mice, but not in Bcl2KO mice when compared to WT mice (Supplemental Fig 2C). In both A1DKO and A1TKO mice, A1a and A1d are lost in all cells raising the possibility that loss of A1 in other cells could have contributed to the decrease in GP66-specific CD4+ T cells. To test this, LCMV-specific SMARTA TCR transgenic CD4+T cells were transferred into A1DKO mice and WT mice and their cell numbers assessed at day 10 after infection. SMARTA T cells expanded similarly in both WT and A1DKO recipient mice (Supplemental fig 2D), indicating that any effects of A1 on other cell types except on T cell survival were negligible.

**Fig 2.**
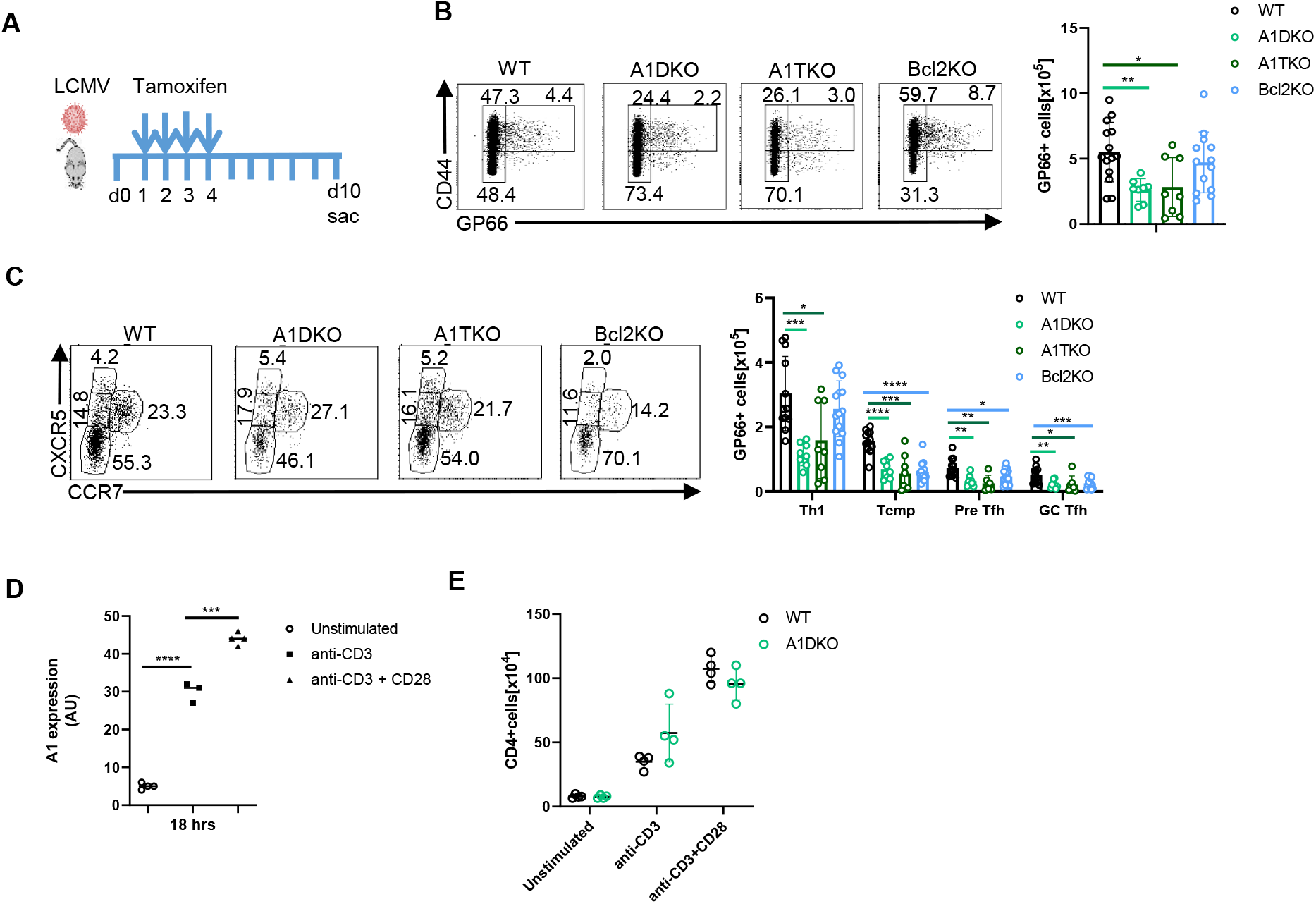
Subset-specific roles for Bcl-2 and A1 in CD4+ T effector survival. (A) Schematic of the experiment for 2B-2C (B) GP66-specifc CD4+ splenic T cells at d10 p.i. with LCMV (C)Within tetramer+ cells, Th1, Tcmp, GC Tfh and pre-Tfh subsets defined by the expression of CXCR5 and CCR7. Numbers in representative plots show the frequency and bar graphs show the numbers in WT (open circle) A1DKO (light green), A1TKO (dark green) and Bcl2KO (blue) mice (D) CD4+ naïve T cells magnetically enriched from spleens of WT mice and cultured without stimulation(open circles) or stimulation with anti-CD3(1 μg/ml, plate-bound) alone(squares) or in combination with anti-CD28(1 μg/ml) antibodies (triangles) for 18 hrs. Cells were harvested, RNA extracted and qPCR was performed for A1 expression. AU= arbitrary units as compared to β-actin. (E) CD4+ T naïve cells magnetically enriched from the spleens of WT(open circle) and A1DKO(light green) mice were activated *in vitro* with anti-CD3 alone (1 μg/mL,plate-bound) or in combination with anti-CD28 antibodies (1 μg/mL) and live cell numbers were estimated after 4 days. Results are representative of at least 3 independent experiments with n=3 or more mice per group and show mean ± SD. \**p* < 0.05, \*\**p* < 0.01, \*\*\**p* < 0.001, **** *p* < 0.0001 Student’s t test.

When broken down into subsets, Bcl2KO mice had similar numbers of Th1 cells but significantly lower numbers of Tcmp and Tfh cells compared to WT mice (Fig 2C). In contrast, the loss of A1 affected the survival of all effector subsets (Fig 2C). Interestingly, no significant differences were observed between A1DKO and A1TKO mice suggesting a minimal role for A1b.

We next determined whether TCR stimulation promoted A1 expression by stimulating CD4+ naïve T cells from WT mice with anti-CD3 and anti-CD28 antibodies *in vitro* and analyzing A1 expression by quantitative PCR. A1 was upregulated by both CD3 and a combination of CD3 and CD28 stimulation (Fig 2D). On the other hand, Bcl-2 was downregulated after CD3 and CD3-CD28 stimulation and Bcl-xL was upregulated only by a combination of CD3 and CD28 stimulation (Supplemental fig 2E)(17). However, TCR-induced A1 expression did not contribute to a survival advantage for activated CD4+ T cells *in vitro* (Fig 2E). Thus, our data show a critical role *in vivo* for A1 in the survival of all viral-specific CD4+ effectors whereas Bcl-2 is important for Tcmp and Tfh survival, but not Th1 survival.

### A1 is vital to control encephalitogenic Th17 CD4+ T cells that drive experimental autoimmune encephalomyelitis (EAE)

As A1 was critical for the survival of expanding CD4+ T effectors, we next determined whether this would impact the development of CD4+ T cell-mediated autoimmunity. We chose a mouse model of multiple sclerosis, because this model has defined peptide antigens with which to track antigen-specific T cells, and it has a demonstrated role for Th17 cells, a subset not observed in LCMV infections. We induced EAE in groups of female A1TKO and WT (C57BL/6) control mice using the standard MOG_35-55_ peptide in CFA protocol (37, 38). By day 12, all the WT controls showed signs of pathogenesis with disease peaking on day 19, however, only one of the A1TKO mice showed signs of pathogenesis which was blunted and did not appear until day 16 (Figure 3A). To assess the role that A1 deficiency had on the development and expansion of MOG-specific T cells we determined the percentage of MOG_35-55_ I-Ab tetramer+ CD4+ T cells in the spleen on day15 post-induction. As can be appreciated in Figure 3B, the percentage of MOG-specific CD4+ T cells in the A1TKO mice were roughly half that in the wild type controls. To determine how A1 deficiency affected EAE-linked cytokines(IL17A, IFNγ, and GM-CSF), we treated mice with Brefeldin A for 4 hours prior to harvesting splenocytes to measure accumulated intracellular cytokines (ICC). A1TKO mice showed a marked reduction in both IL-17A as well as IL-17A and IFNγ double producing splenic CD4+ T cells compared to controls (Figure 3B). Taken together, these data strongly suggest that CD4+ T cells lacking A1 at the time of EAE induction develop markedly reduced disease with an accompanying reduction in MOG-specific Th17 cells and pathogenic cytokine production.

**Figure 3.**
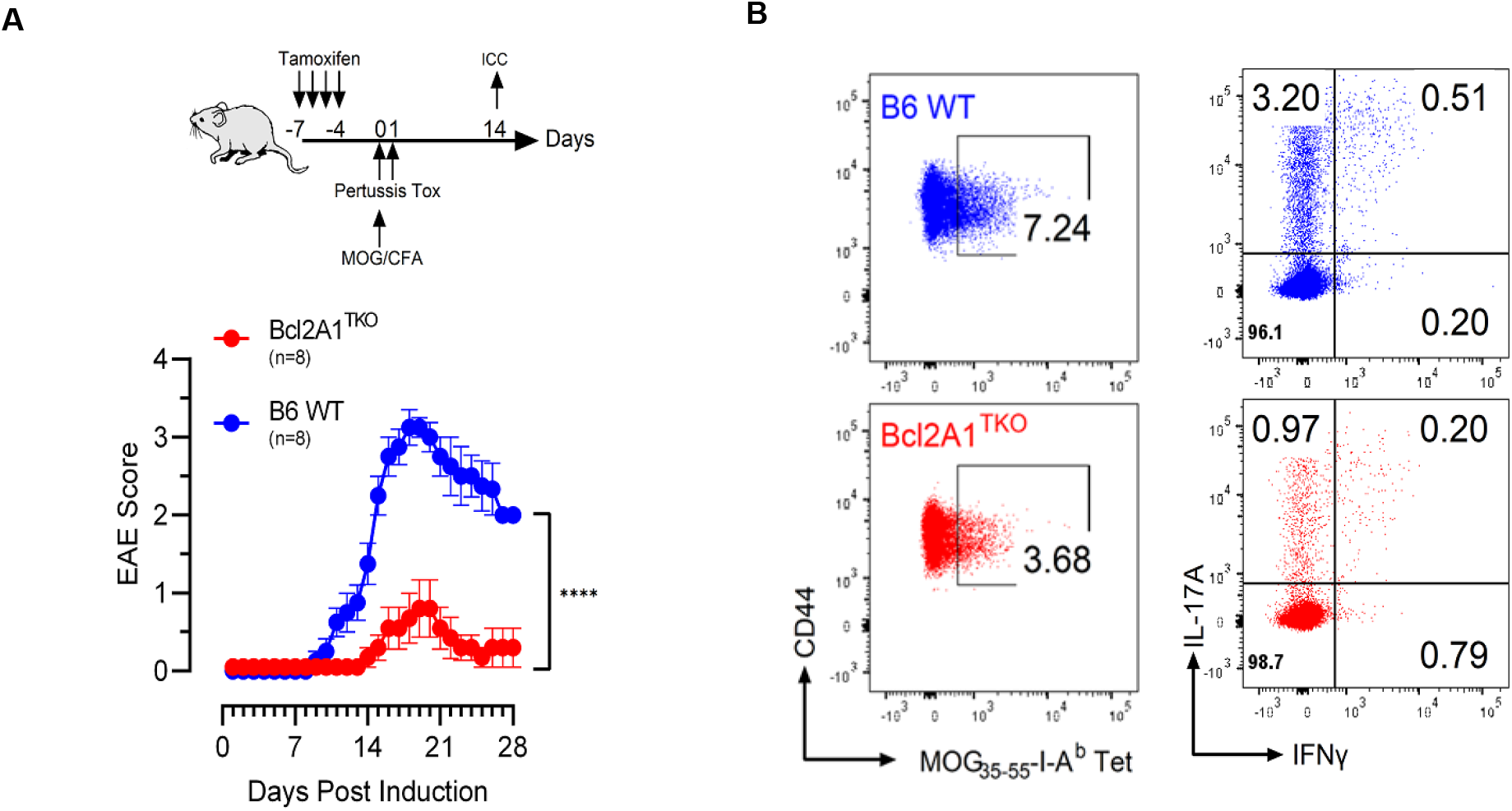
A1TKO mice have reduced EAE and decreased levels of MOG-specific Th17 cells with reduced inflammatory cytokine production. A1TKO mice were treated with tamoxifen for 4 days to induce the loss of the A1b gene, then rested for 3 days (A)The resulting A1TKO mice and B6 WT controls were injected with MOG_35-55_ peptide (100µg) in CFA (Hooke Laboratories) and treated on Day 0 and +1 with 80ng of pertussis toxin to induce EAE. Mice were then followed daily for signs of EAE. Reduced EAE development is seen in A1TKO as compared to controls. All eight female A1TKO mice showed significantly reduced EAE (****; p≤0.001, Pearson two-tail), based on the scoring system described in the methods section (B)Reduced MOG_35-55_-I-A_b_ tetramer_+_ T cells in A1TKO mice with reduced levels of IL-17A, IFNγ and IL-17A/IFNγ double producing CD4_+_ T cells. On Day 14, one A1TKO and one WT mouse, randomly selected, were treated with Brefeldin A for 4 hours prior to sacrifice and splenocytes were harvested for analysis by flow cytometry. T cells were gated on size, viability, CD4_+_, CD44_+_ and either MOG-tet expression or for intracellular cytokines (IL-17A, IFNγ) assessed. Data are cumulative of 2 independent experiments.

### Bcl-2 is critical to antagonize Bim and promote the CD4+ effector to memory transition

After their proliferative expansion, most effector CD4+ T cells die via Bim-dependent apoptosis, while some survive and become memory T cells. The mechanism(s) by which individual effector CD4+ T cell subsets restrain Bim and survive as memory T cells remains unclear. To determine the expression of Bcl-2 family members in surviving GP66-specific CD4+ T cells, we sorted these cells from the spleen of WT mice on day 24 after LCMV infection and subjected them to sc-RNAseq. By day 24, the GP66-specific CD4+ T cell subsets exhibited dynamic alterations in pro- and anti-apoptotic Bcl-2 family member expression; Bim and Bcl2 were increased in Th1 and Tcmp subsets compared to A1b and A1d which decreased (Supplemental Fig 3A). Further, both sc-RNAseq and flow cytometric analysis revealed that although most subsets remained post contraction, the frequency of individual subsets changed, with Th1 cells being reduced and Tcmp cells being increased in frequency (Fig 4A). When we assessed the absolute numbers of LCMV-specific T cells, Tcmp cells contracted less (∼ 50%) from d10 to d21 whereas most of the Th1 cells were lost (∼90%) during that time (Fig 4B). We next assessed the Bcl-2/Bim ratios in Tcmp and Th1 cells at d10 p.i. Flow cytometric analysis showed that Tcmp cells had a significantly increased Bcl-2/Bim ratio at the peak of the response compared to Th1 cells (Fig 4C). Further, Bcl-2 was significantly increased in Th1 cells that remained after day 20 post infection in comparison to Th1 cells at day 10 (Fig 4C), suggesting that cells with the highest Bcl-2 expression were selected to survive.

**Fig 4.**
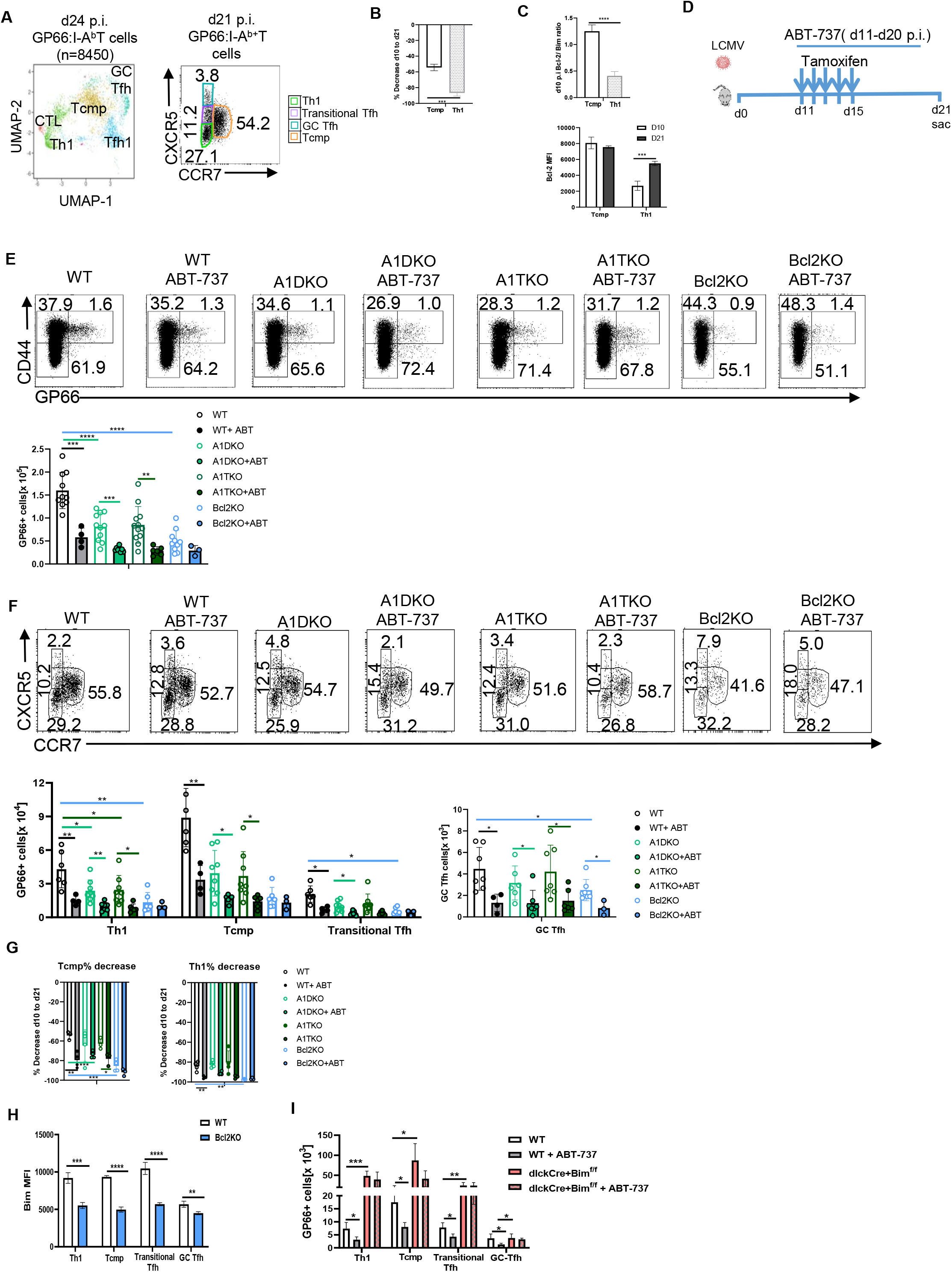
Bcl-2 is critical to counteract Bim and promote the CD4+ T cell survival during contraction. (A) sc-RNAseq of splenic GP66-specific CD4+ T cells from WT mice at d24 p.i. with LCMV. UMAP plot showing clusters identified by variable gene expression cells in four mice at day 24 p.i. Each dot is an individual CD4+ T cell and the colors highlight clusters. Flow cytometric analysis of GP66-specific CD4+ splenic T cells at d21 p.i with LCMV defined as Th1, Tcmp, GC Tfh and transitional Tfh subsets by the expression of CXCR5 and CCR7 (B) Percent decrease of Th1 and Tcmp cells from d10 to d21 p.i. in WT mice (C) Bcl-2/Bim ratio in Tcmp vs Th1 at d10 p.i. (top) and Bcl-2 expression in Th1 and Tcmp at d10 and d21p.i. in WT mice (bottom). (D) Schematic of the experiment for 4E-4G (E) GP66-specific CD4+ splenic T cells at d21 p.i. with LCMV (F) Within tetramer+ cells, representative plots of frequency and bar graphs of cell numbers of Th1, Tcmp, Tfh and transitional Tfh subsets defined by the expression of CXCR5 and CCR7(G) Bar graphs of the percentage decrease of Th1 and Tcmp cells from 10 to d21 p.i. in WT(open circles), WT + ABT-737(black circles), A1DKO (light green open circles), A1DKO+ABT-737(light green filled circles), A1TKO (dark green open circles), A1TKO+ABT-737(dark green filled circles),Bcl2KO(open blue circles) and Bcl2KO + ABT-737 (filled blue circles) mice. (H) Bim MFI in WT (white) and Bcl2KO (blue) mice in GP66-specific CD4+ Th1, Tcmp, GC Tfh and transitional Tfh subsets (I) Cell numbers of GP66-specific CD4+ T cells at d21 post infection with LCMV in WT mice(open), WT+ABT-737(grey), dlckCre+Bim_f/f_ (peach) and dlckCre+Bim_f/f_ + ABT-737(peach+grey) mice. Results are representative of at least 3 independent experiments with n=3 or more mice per group and show mean ± SD. \**p* < 0.05, \*\**p* < 0.01, \*\*\**p* < 0.001, **** *p* < 0.0001 Student’s t test.

To formally test the role of Bcl-2 in the CD4 effector to memory transition we treated Bcl2KO mice with tamoxifen from d11 to d15 after LCMV infection (Fig 4D) to delete Bcl-2 during the contraction phase of the response (Supplemental Fig 3A). Additionally, to test for potential redundancy between Bcl-2, A1, and Bcl-xL, we treated WT, A1DKO, A1TKO and Bcl2KO mice with ABT-737 to inhibit Bcl-2 and Bcl-xL from day 11-20 after LCMV infection (Fig 4D). Both genetic ablation (Bcl2KO) and pharmacologic inhibition of Bcl-2 (ABT-737 treatment) resulted in a significant loss of total GP66-specific CD4+ T cells (Fig 4E). Specifically, Bcl-2 was critical for survival of Tcmp cells, with Bcl2KO mice having a ∼4 fold loss compared to WT mice (Fig 4F), which translated to an ∼80% decrease from d10 to d21 p.i. compared to a 50% decrease in WT mice (Fig 4G). Timed loss of Bcl-2 also significantly impacted Th1 survival leading to ∼2 fold loss (Fig 4F) and a ∼ 95% decrease in Th1 cells (Fig 4G). By d21 after LCMV infection, most GC Tfh cells downregulate CXCR5 expression (13), hence CXCR5^int^ CCR7-cells are likely a mix of cells that haven’t fully differentiated into GC Tfh(39) and Tfh cells with downregulated CXCR5 levels. Therefore we refer to these CXCR5^int^ CCR7-cells as transitional Tfh. Both transitional Tfh and GC Tfh cells were significantly impacted by the loss of Bcl-2 (Fig 4F). Treatment of WT, A1DKO, A1TKO mice with ABT-737 decreased the absolute numbers of all subsets of GP66-specific CD4+ T cells to levels observed in Bcl2KO mice (Fig. 4E, F). Although ABT-737 also inhibits Bcl-xL (and Bcl-w) we did not see an additional loss of GP66-specific CD4+ T cells in ABT-737-treated Bcl2KO mice in comparison to Bcl2KO mice, with the exception of GC Tfh cells (Fig 4F). Together, these data suggest, in most CD4+ T cell subsets, the effects of ABT-737 are largely due to its inhibition of Bcl-2 rather than potential redundant role for Bcl-xL.

Although the numbers of GP66-specific CD4+ T cells was decreased in A1DKO and A1TKO mice on d21 after infection (Fig 4E, 4F), we note that these mice had already decreased numbers of GP66-specific CD4+ T cells at the peak of the response (Fig 2B,2C). To account for this, we calculated the decline of Th1 and Tcmp subsets from d10 to d21 p.i. and found that the relative loss of cells in A1DKO and A1TKO mice was not different from the loss in WT mice (Fig 4G). Further, the numbers of GP66-specific CD4+ T cells were not significantly different between ABT-737 treated A1TKO and A1DKO in comparison to Bcl2KO mice (Fig 4E), indicating that Bcl-2 was the major driver of survival during contraction of the response.

As Bim is critical for contraction of CD4+ T cell responses, we assessed Bim levels in GP66-specific CD4+ T cells on d21 p.i. (after inducible deletion of Bcl-2 from d11 to d15) and saw that cells that had lost Bcl-2 had significantly lower levels of Bim (Fig 4H). To determine whether Bcl-2 antagonized Bim to promote survival of GP66-specific CD4+ T cells, we treated WT and Bim-deficient mice with ABT-737. In WT mice, treatment with ABT-737 from d11 to d20 after infection significantly reduced the survival of all subsets of GP66-specific CD4+ T cells (Fig 4I). However, in mice with a T cell specific loss of Bim, dLckCreBim^fl/fl^ (40), ABT-737 failed to drive the death of all subsets of GP66-specific CD4+ T cells (Fig 4I). Taken together, Bcl-2 is critical for antagonizing Bim in all subsets of GP66-specific CD4+ T cells during contraction of the response.

### IL-7 and IL-15 drive the survival of Th1 cells during contraction

IL-7 and IL-15 are critical to maintain Bcl-2 expression during contraction of the CD8+ T cell response (5, 6, 34), however their role in CD4+ T cells is unclear. Expression of IL-7Rα (CD127) on GP66-specific CD4+ T cells was dynamic, having low expression on day 7 which was substantially upregulated by day 21 after infection (Fig 5A). In contrast, CD122 (the IL-2Rβ chain, shared by IL-2 and IL-15 signaling) was expressed on some cells in all subsets on day 7 but was upregulated on cells in all subsets by day 21 (Fig 5A). Flow cytometric analysis at d21 post infection in WT mice showed that CD127 was expressed on most splenic GP66-specific Th1 and Tcmp cells but hardly expressed on GC-Tfh cells, while CD122 was highest on Th1 cells and transitional Tfh, but low on Tcmp and GC Tfh cells(Fig 5B). Further, GP66-specific CD127+ cells significantly higher levels of Bcl-2 than their CD127-counterparts across all subsets (Fig 5C top). Although, Bcl-2 levels were not different between CD122+ and CD122-in GP66-specific CD4+ T cells, Bcl-2 levels were significantly increased in CD122+ Th1 and transitional Tfh subsets relative to their CD122-counterparts (Fig 5C bottom). Though our prior work revealed a minimal role for IL-7 and IL-15 in memory CD4+ T cell survival after contraction (6, 41), subset-specific roles were not examined. To determine the individual and combined roles of IL-7 and/or IL-15 on GP66-specific CD4+ T cell survival, we treated WT mice with neutralizing antibodies for IL-7 and/or IL-15. The combined, but not individual, neutralization of IL-7 and IL-15 led to a significant loss in the frequency and numbers of Th1 cells at day 21 of the response, whereas other subsets remained unaffected (Fig 5D). Together, these data are consistent with a scenario in which, during contraction of the response, IL-7 and IL-15 promote expression of Bcl-2 in Th1 cells and are important for their survival.

**Fig 5.**
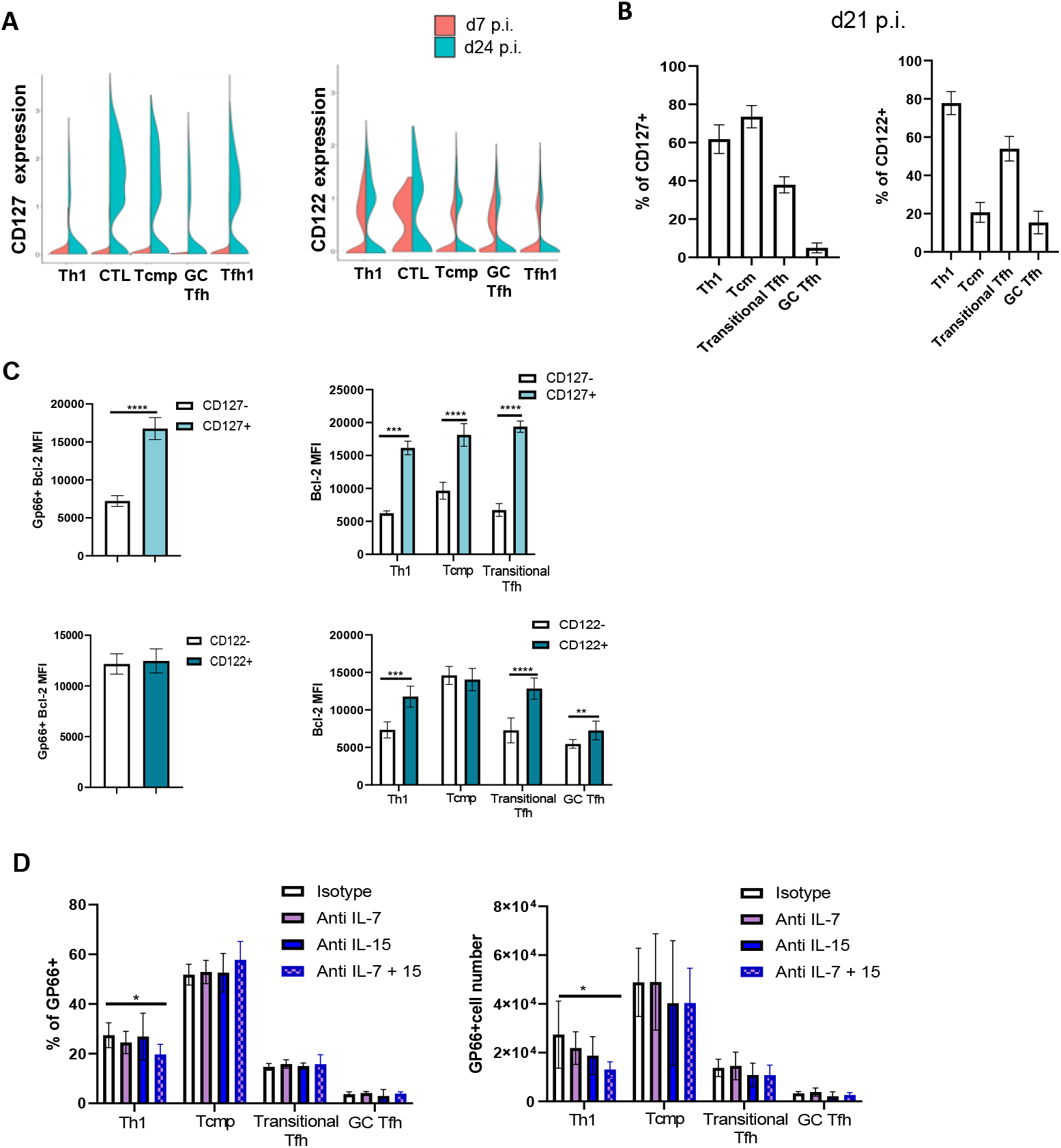
IL-7 and IL-15 driven survival of Th1 cells during contraction. (A) CD127 and CD122 mRNA expression from scRNAseq of GP66-specific CD4+ T cell clusters in WT mice at d7 versus d24 post LCMV infection (B) Flow cytometric analysis of frequency of CD127 and CD122 expressing cells in GP66-specific CD4+ T subsets at d21p.i. with LCMV in WT mice (C) Bcl-2 expression in CD127-vs CD127+ GP66-specific CD4+ T cells and in CD127-vs CD127+ Th1, Tcmp and transitional Tfh subsets within GP66-specific CD4+ T cells at d21 p.i. in WT mice. GC Tfh cells that expressed CD127 were negligible and hence an MFI of Bcl-2 could not be determined(D) Bcl-2 expression in CD122-vs CD122+ GP66-specific CD4+ T cells and in CD122-vs CD122+ Th1, Tcmp, transitional Tfh and GC Tfh subsets within GP66-specific CD4+ T cells at d21 p.i. in WT mice (E) d21 p.i. frequency and cell numbers of GP66-specific CD4+ T cell subsets in WT mice treated intraperitoneally with 3 mg of either isotype control(white), anti IL-7 (M25)(purple), anti-IL-15(M96)(dark blue) or both anti-IL-7 and IL-15(blue and purple) on days 11,13,15,17 and 19 p.i. with LCMV. Results are representative of 2 independent experiments with n=3 or more mice per group and show mean ± SD. \**p* < 0.05, \*\**p* < 0.01, \*\*\**p* < 0.001, **** *p* < 0.0001 Student’s t test.

### Bcl-2 is critical for the generation and survival of memory CD4+ T cells

During contraction of the CD4+ T cell response, some cells survive and become memory cells. We next determined whether Bcl-2 expression during this timeframe was critical for development of long-term memory. Strikingly, tamoxifen-driven loss of Bcl-2 during contraction of the response (Fig 6A) resulted in significantly fewer GP66-specific CD4+ memory T cells 90 days after infection (Fig 6B). At this late time point, the subsets were comprised predominantly of Th1 cells (CXCR5-CCR7-, 45%), Tcm cells (CXCR5^int^ CCR7+, 30%), transitional Tfh cells (CXCR5^int^ CCR7-, 25%) with very low numbers of GC Tfh cells (CXCR5+CCR7-), consistent with prior work (10) and all subsets required Bcl-2 during contraction of the response for their long-term survival (Fig 6C). Functionally, memory CD4+ T cells are endowed with an ability to rapidly produce multiple cytokines (42, 43). To determine whether Bcl-2 altered the numbers of cytokine-producing memory CD4+ T cells, we assessed their ability to produce IL-2, TNFα and IFNγ after brief stimulation of splenocytes with GP66-80 peptide *in vitro* and intracellular staining for cytokine production. Strikingly, there was a ∼ 5 fold loss in frequency and cell numbers of IFNγ, TNFα and IL-2 producing cells at d90 p.i. in mice with Bcl-2 deletion during contraction, compared to WT mice (Fig 6D). Similarly, pharmacologic inhibition of Bcl-2 function with ABT-737 from d11 to d20 after LCMV infection also resulted in a significant loss in frequency and numbers of IFNγ, TNFα and IL-2 producing cells 150 days later (Fig 6E). Combined, these data show that Bcl-2 is required during contraction of the response for the long-term survival and generation of functional memory CD4+ T cells.

**Fig 6.**
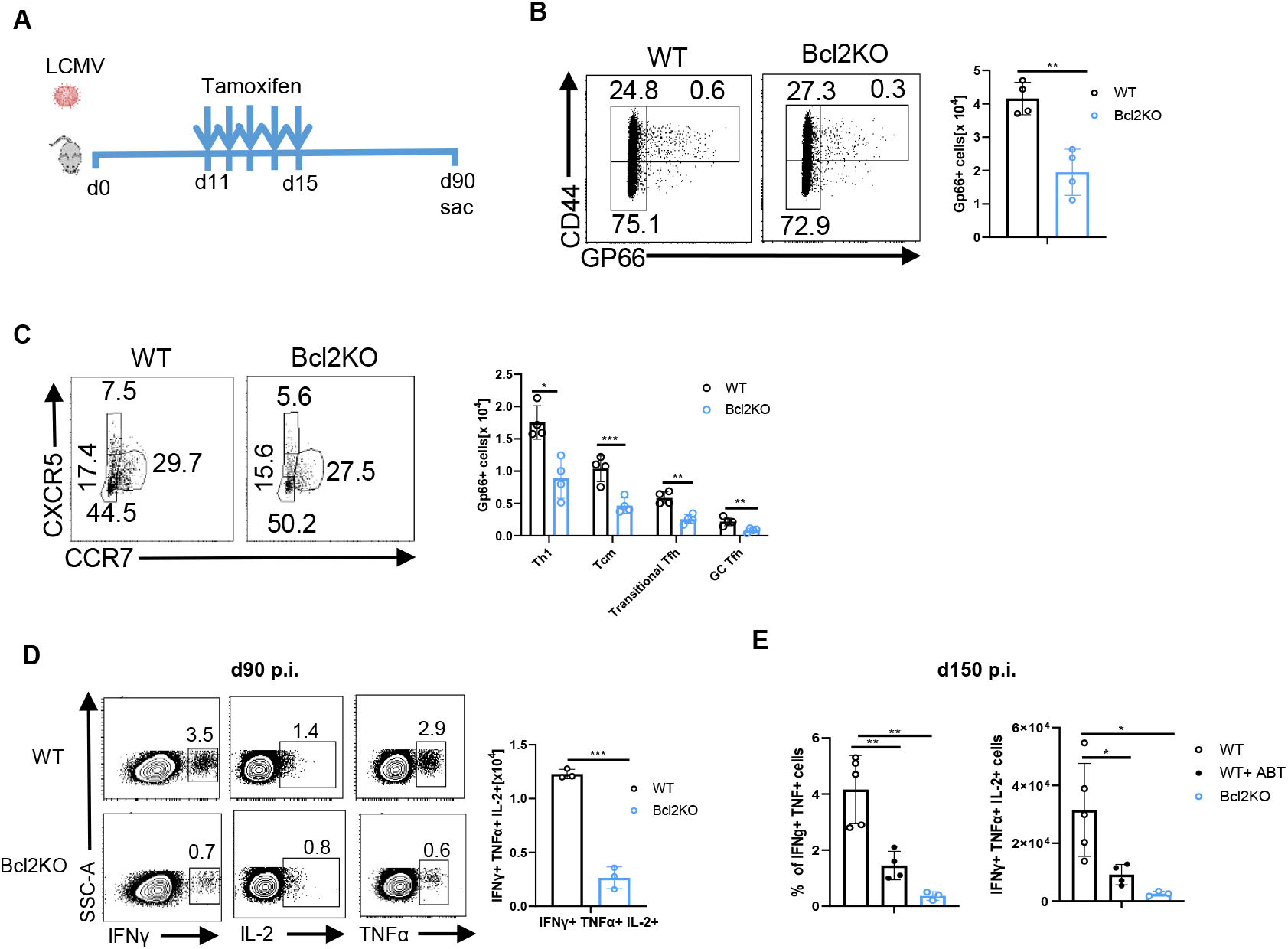
Bcl-2 is required during contraction for the generation of long-lived memory CD4+ T cells. (A) Schematic of experimental procedure for B-D (B) GP66-specific CD4+ splenic T cells at d90 p.i. with LCMV (C) Th1, Tcm, GC Tfh and transitional Tfh subsets within GP66-specific CD4+ T cells defined by the expression of CXCR5 and CCR7 at d90 p.i. (D) IL-2, TNFα and IFNγ positive cells at d90 p.i. Representative plots show frequency and bar graphs show cell numbers in WT (white) and Bcl2KO (blue) mice when Bcl-2 was deleted at d11 to d15 p.i (E) Cell numbers of IL-2, TNFα and IFNγ triple producers at d150 p.i. in WT(open circle), inhibition of Bcl-2 function by treatment with ABT-737 from d11 to d20 p.i.(black filled circles) and tamoxifen-mediated deletion of Bcl-2 from d11 to d15 p.i.(blue circles).Results are representative of 2 independent experiments with n=3 or more mice per group and show mean ± SD. \**p* < 0.05, \*\**p* < 0.01

### Bcl-2 is critical for long-term maintenance of memory CD4+ T cells

In addition to its role during contraction of the response, Bcl-2 is also likely critical to maintain memory CD4+ T cell survival after their formation, potentially acting downstream of IL-7 (44-47). To this end, we temporally deleted Bcl-2 from days 85 to 89 after infection and analyzed GP66-specific CD4+ T cells a day after the last tamoxifen injection (Fig 7A). Deletion of Bcl-2 in CD4 memory cells, resulted in a 3-fold reduction of GP66-specific CD4+ T cells compared to WT mice (Fig 7B) with all memory subsets being significantly affected (Fig 7C). Thus, Bcl-2 is critical for the maintenance of viral-specific memory CD4+ T cells.

**Fig 7.**
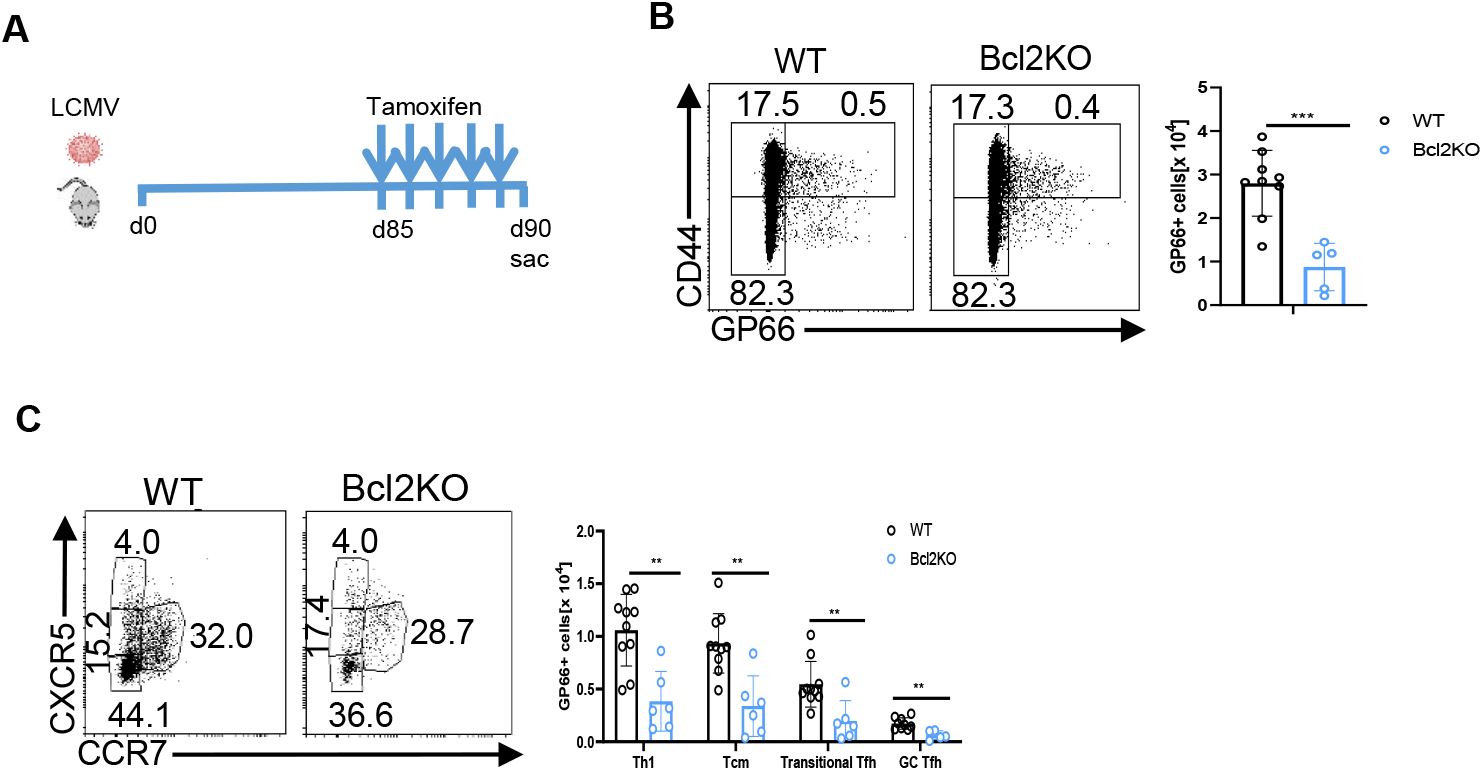
Bcl-2 is critical for long term maintenance of memory CD4+ T cells. (A) Schematic of experimental procedure (B) GP66-specific CD4+ splenic T cells at d90 p.i. with LCMV(C) Th1, Tcm, GC Tfh and transitional Tfh subsets within GP66-specific CD4+ T cells defined by the expression of CXCR5 and CCR7 at d90 p.i. Representative plots show frequency and bar graphs show cell numbers in WT (white) and Bcl2KO (blue) mice in which Bcl-2 was deleted at d85 to d89 p.i. Results are representative of 2 independent experiments with n=3 or more mice per group and show mean ± SD. \**p* < 0.05, \*\**p* < 0.01

## Discussion

The experiments outlined in this paper represent our continuing efforts to understand the role of individual Bcl-2 family members in control of T cell homeostasis (2, 3, 5-7, 34, 41, 48-50). Here we report novel temporal and subset specific roles for A1 and Bcl-2 in the survival of CD4+ T cells responding to viral infection. Interestingly, we found that A1 is required for the survival of most effector CD4+ T cell subsets during expansion, including the largest Th1 subset, but also smaller subsets of Tfh and Tcmp cells. Similarly, we found that Th17 cells induced by immunization also relied on A1 and that absence of A1 significantly reduced susceptibility to EAE disease. Thus, A1 is critical for the generation of normal numbers of effector CD4+ T cells amongst all subsets measured thus far. Given these data, we think that the requirement for A1 occurs early, after their initial TCR stimulation, but before differentiation potentially as part of a gene expression module enacted to promote survival during proliferation. Alternatively, the requirement for A1 could manifest equally across effector subsets after their differentiation. Future experiments are underway to examine the requirement for A1 before and after subset differentiation.

A recent report showed that mice deficient in A1a/b/d did not have lower numbers of CD4+ T cells after influenza infection (19), however there were differences between the two studies, including the strains of viruses and infectious doses used. When we examined a small fraction of antigen-specific CD4+ T cells during LCMV acute infection, we found that they were significantly decreased. We found similar results after MOG peptide immunization during EAE. Thus, further work will be needed to determine whether the requirement for A1 is model-specific. Our data also show that, although there are 3 functional A1 genes, it appears that A1a and A1d are critical as there was no difference in the numbers of effector CD4+ T cells in the presence or absence of A1b suggesting differential roles for the 3 A1 genes despite their high homology. Prior work showed that A1b is highly expressed in resting T cells while A1a and A1d are induced upon activation (51). Finally, as not all Th1 cells were lost in the absence of A1, it is likely that additional anti-apoptotic Bcl-2 family members contribute redundantly to A1. A likely candidate is Bcl-xL, given that T cell activation and co-stimulation upregulate the levels of Bcl-xL.

Although initial work suggested that CD28-driven expression of the anti-apoptotic protein, Bcl-xL is important for activated T cell survival, these experiments were performed *in vitro* (52). However, *in vivo*, T cell-specific loss of Bcl-xL, did not impair T cell responses to acute bacterial infection (9). Unlike Bcl-xL, we found A1 to be critical for the survival of antigen-specific effector CD4+ T cells during expansion *in vivo*. Although we and others have found that TCR and CD28 signaling can induce expression of A1 *in vitro*, they are likely not the sole factors controlling A1 expression *in vivo*. For example, our sc-RNA seq at day 7 post-infection revealed that expression of A1b and A1d, was correlated with the co-stimulatory molecules OX40 and CD30L (data not shown). As OX40 is induced after CD28 co-stimulation (53) and controls effector CD4+ T cell survival (54), it is possible that CD28, OX40 and CD30L contribute individually or redundantly to A1 expression during proliferative expansion of effector CD4+ T cells. Further work is required to determine the individual roles of CD28, CD30L and OX40 on induction of A1 in effector CD4+ T cells.

Given the role for A1 in CD4 effector survival, the pro-apoptotic molecules(s) that are antagonized by A1 remain unclear. A1 is known to bind Bim, Puma and Noxa (55) and there could be subset-specific binding partners at play. For example, GC Tfh cells that depend on late TCR signals (56), expressed high levels of A1, low Bim and high Noxa. As A1 is a high affinity binding partner for Noxa (57), A1 could be counteracting Noxa in these cells. Conversely, Th1 and Tcmp, expressed high levels of Bim and very little Noxa, so A1 could be antagonizing Bim in these subsets during the expansion phase of the response.

At the peak of the response, Bcl-2 levels are significantly lower in Th1 cells and Bcl-2 is not required for their survival during proliferative expansion. On the other hand, Tfh and Tcmp cells have high levels of Bcl-2 at the peak of the response, and require Bcl-2 for their survival, similar to naïve CD4+ T cells (34).One explanation for this result is that Bcl-2 is selectively downregulated in Th1 cells, but maintained in Tfh and Tcmp cells. Alternatively, it could be that Bcl-2 is transcriptionally repressed in all effector CD4+ T cells, but selectively upregulated in Tfh and Tcmp cells. In favor of the former, recent data suggests that high TCR signal strength drives Th1 differentiation (58-60) and manipulation of TCR signal strength by stimulation with greater amounts of peptide led to decreased Bcl-2 levels(17). Thus, Tfh and Tcmp cells could be experiencing lower TCR stimulation and hence be able to maintain high levels of Bcl-2 that prolong their survival. Taken together, during proliferative expansion, strong TCR signals induce a programmatic switch in Bcl-2 family members that controls CD4+ T cell survival.

After the peak of the response, effector Th1 and Tcmp switch away from an A1-mediated survival to a Bcl-2-mediated survival. The short half-life of A1(61, 62),the lack of antigen that drives TCR and/or co-stimulatory signals promoting A1 expression and the low levels of Bcl-2 could render most effector CD4+ T cells susceptible to death during contraction of the response. However, some Th1 cells and most Tcmp cells express high levels of Bcl-2 and these cells are significantly enriched during contraction of the response. Additionally, the propensity for contraction varies among distinct CD4+ T cell subsets, with Tcmp dying significantly less than Th1. Notably, our data strongly suggest that the high Bcl-2/Bim ratio in Tcmp afford these cells a significant protection from death during contraction compared to Th1 cells. The upstream regulators of Bcl-2 in Tcmp require further investigation as the combined neutralization of IL-7 and IL-15 only affected Th1 survival. Nonetheless, after the peak of the response, effector CD4+ T cells switch back from a TCR-driven survival program involving A1 to a Bcl-2-mediated survival program to form memory. Strikingly, the loss of Bcl-2 function during contraction of the response led to a long-term loss of all memory CD4+ subsets and an even more dramatic loss in cytokine-producing viral-specific CD4 T cells. Interestingly, the loss of Bcl-2 long after contraction and during the memory maintenance phase, also led to a loss of viral-specific memory cells, but did not significantly affect their cytokine production(Supplemental Fig 4). Thus, our data suggest that during contraction of the response, a CD4+ memory precursor subset that matures into full-fledged, poly-cytokine producing memory cells requires Bcl-2 for its survival. Further work is underway to determine the cytokine-producing potential of CD4 effector and memory T cell subsets and their dependence on Bcl-2.

Taken together, our data clearly show that effector CD4+ T cells display a dynamic dependence upon Bcl-2 family members for survival as they progress from activation to memory formation. This knowledge could be exploited to manipulate subset-specific CD4+ T cell survival during distinct phases of the response. For instance, the high dependence of activated T cells on A1 could be exploited by use of A1 inhibitors to target unwanted T cells during early onset disease, as our data suggest would be efficacious in EAE. On the other hand, Bcl-2 inhibitors, such as venetoclax, might be employed to reduce pathogenic memory T cells that promote auto-, allo-immunity or allergy, without diminishing Th1 cells responsive to infection. Therefore, Bcl-2 family member antagonists hold promise to target pathogenic CD4+ T cells in the context of transplantation, autoimmune and allergic diseases. Conversely, knowledge of factors driving subset-specific CD4+ T cell survival can be used to boost immunity, for instance in optimal vaccine design.

## Acknowledgements

We thank the members of the Hildeman, Chougnet, and Jordan labs for helpful discussion. This research was also made possible, in part, using the Cincinnati Children’s Single Cell Genomics Core [RRID: SCR_022653], DNA Sequencing and Genotyping Core [RRID: SCR_022630], and Biomedical Informatics Core. We specifically acknowledge the assistance of Kelly Rangel and Shawn Smith from the Single Cell Genomics Core. This work was supported by Public Health Service grants AI142264 (D.A.H, E.S.W.), AI057753 (D.H.). All flow cytometric data were acquired using equipment maintained by the Research Flow Cytometry Core in the Division of Rheumatology at Cincinnati Children’s Hospital Medical Center which is supported in part by NIH AR070549 and by the Hematology Center of Excellence grant U54DK126108.

## Materials and methods

### Mice

C57BL/6 mice were purchased from Taconic Farms. Bcl2^f/f^ mice, a gift from Dr. Ira Tabas (Columbia University, New York, NY) and A1TKO mice, a gift from Dr Marco Herold (36) were crossed to *Cd4-creER*^*T2*^ mice obtained from The Jackson Laboratory. Bim^f/f^ mice were generated at the Walter and Eliza Hall institute in collaboration with Dr. Philippe Bouillet as previously described(63) and were crossed to dLckCre mice obtained from The Jackson Laboratory. Mice between 8 to 12 weeks of age were used for infections. Experimental and control mice were littermates in most cases, and both male and female mice were used. Wild-type (WT) mice mentioned in the figure legends are C57BL/6 or *Cd4-creER*^*T2-*^ Bcl2^f/f^ mice; we have not observed significant differences between them. Animals were housed under specific pathogen-free conditions in the Division of Veterinary Services (Cincinnati Children’s Hospital Medical Center, Cincinnati, OH), and experimental procedures were reviewed and approved by the Institutional Animal Care and Use Committee at the Cincinnati Children’s Hospital Research Foundation.

### Infection model and tamoxifen administration

Mice (8-12 weeks old) were infected by intra peritoneal injection with 2 × 10^5^ pfu of LCMV Armstrong. In experiments using *Cd4-creER*^*T2*^ animals, tamoxifen (2mg/mouse/day/100 ul volume) or corn oil controls were injected by intra peritoneal injection at the indicated time points.

### T cell stimulation and qRT-PCR

CD4+ T naïve T cells from the spleens of WT mice were isolated by negative selection using the EasySep Mouse Naïve CD4^+^ T cell Isolation kit (StemCell Technologies). T cells were activated with 1 *μ*g/ml plate-bound anti-CD3 (BioLegend, clone 145-2C11) and 1 *μ*g/mL anti-CD28 (BioLegend, clone 37.51) antibodies for 18 hrs. Cells were processed for RNA (RNeasy micro kit, Qiagen) and reverse transcribed to cDNA (SuperScript First Strand Synthesis Kit for RT-qPCR, Invitrogen) and qPCR was carried out (TaqMan Universal PCR Master Mix, no AmpErase^™^UNG and Applied Biosystems StepOnePlus cyler). TaqMan Probes (ThermoFisher) A1 (Mm03646861_mH), Bcl-xL (Mm00437783_m1), Bcl-2(Mm00477631_m1) and Actb(Mm02619580_g1) and the following primers were used: A1 fwd 5’ TTTCCAGTTTTGTGGCAGAAT 3’; A1rev 5’ TCAAACTTCTTTATGAAGCCATCTT 3’; Bcl-2 fwd 5’ AGTACCTGAACCGGCATCTG 3’; Bcl-2 rev 5’ AGGGTCTTCAGAGACAGCCA 3’; Bcl-xl fwd 5’ GTTGGATGGCCACCTATCTG 3’; Bcl-Xl rev 5’ GCTGCATTGTTCCCGTAGAG 3’. PCR conditions: 10 min 95°C; 40 cycles x (15 sec 95°C; 1 min 60°C) Relative quantification was calculated using the ΔΔCT method.

### Cell processes and flow cytometry

Spleens from individual mice were harvested and crushed through a 100 μm mesh strainer to generate single-cell suspensions. Two to three million cells were stained with tetramers that recognize H-2 IA^b^-GP66 tetramer (NIH Tetramer Core Facility) at 37°C for 1 hr 15 min followed by staining with other surface antibodies (except CXCR5) at 37°C for 45 min. After the tetramer and surface stains at 37°C, CXCR5 staining was performed for 30 min at 4°C. Antibodies used were CD4, CD44, TCRβ, CD16/32, (BD Biosciences), CXCR5, CCR7, Bcl-2, TNFα, IFNγ, IL-2 (Biolegend), CD127, CD122, live dead blue viability dye (Invitrogen) and Bim (Cell Signaling Technology). Cytokine staining was performed after 5 hours on splenocytes incubated in the presence of GP_61-80_(GLKGPDIYKGVYQFKSVEFD, 10 μg/ml) peptide in the presence of brefeldin A and monensin for the final 4 hours, stained for surface markers and then followed by intracellular staining. Intracellular stains were performed using the Foxp3/transcription factor staining buffer set (Invitrogen). The cells were acquired on a BD LSRFortessa flow cytometer and analyzed by FACSDiva software (BD Biosciences) or FlowJo software (Tree Star). Cell sorting was performed on a FACSAria II (BD).

### Active EAE

Groups of female WT or A1TKO mice were immunized s.q. with 100 µg MOG_35-55_ emulsified in 5mg/ml CFA (Hooke Laboratories, Lawrence, MA) per manufacturor’s protocol. On days 0 and 1, mice were injected with 80 ng pertussis toxin, i.p. (Hooke Laboratories). On Day 14 post after induction, MOG-specific CD4+ T cells enumerated in the spleen was performed using MOG_35-55_-I-A^b^-PE tetramers (NIH Tetramer Core, Emory University, Atlanta, GA) and a panel of fluorochrome labeled antibodies to CD45 (30-F11-AF647), CD4 (GK1.5-APC), CD16/32 (93-FITC), CD44 (IM7-BV711) to assess Th subset development and activation states. In addition to assess MOG-driven cytokine production, we re-stimulated T cells with MOG_35-55_ peptide pulsed APC for 24h in culture (Brefeldin A treatment last 4-6 hr) and assessed cytokine production by intracellular cytokine staining and flow cytometry for IL-17 (TC11-18H10.1-BV421) and IFNγ (XMG1.2-AF700). The remaining mice (8 mice per group) were monitored twice daily for disease following a clinical score metric based on the following scale: 0, no symptoms; 1, tail flaccidity; 2, hindlimb weakness and/or ataxia; 3, hindlimb paralysis; 4, hindlimb and forelimb paralysis; 5, moribund. Immunization and pertussis toxin injections were formulated to maximize the disease severity to ≤3, per the manufacturer’s instructions and stipulated in our approved IACUC protocol (IACUC 2020-0080).

### In vivo cytokine neutralization

Anti-IL-7(clone M25) and Mouse IgG2b, κ isotype control (clone MPC-11) were purchased from BioXcell. Anti-IL-15 was a kind gift from Amgen (Seattle, WA). For IL-7 neutralization, infected mice were injected with i.p. with 3 mg of anti-IL7 or isotype control on days 11, 13, 15, 17 and 19 post infection, sacrificed at d21 and spleens harvested. For IL-15 neutralization, infected mice were injected i.p. with 25µg of M96 on days 11, 13, 15, 17 and 19 post infection, sacrificed at d21 and spleens harvested. IL-7 neutralization was confirmed by > 80% loss of immature B cells in the bone marrow and IL-15 neutralization was confirmed by >60% loss of splenic natural killer cells (data not shown).

### Single cell RNA-Seq

5000-10000 splenic GP66:I-A^b+^ T cells were sorted from LCMV infected mice, loaded onto a 10X Chromium platform to generate cDNA carrying cell-specific barcodes that were used to construct sequencing libraries using the Chromium Single Cell 3′ Library & Gel Bead Kit v2 according to the manufacturer instructions. Libraries were sequenced and raw base call files were de-multiplexed with Cell Ranger (26) v3.1.0 mkfastq (10x Genomics). Reads were aligned to mouse reference genome mm10 and gene expression quantified using Cell Ranger count. Further data analysis was carried out with Seurat (64, 65) v3.1.3in R v3.5.0 (66).Cells displaying more than 10% mitochondrial gene expression, or less than 500 total expressed genes were excluded from the analysis. Gene expression counts were normalized with the NormalizeData function in Seurat, which uses a logarithmic normalization method where gene counts for each cell are divided by its total counts and natural log-transformed using log1p and multiplied by a scale factor of 10,000. The four samples were integrated together using FindIntegrationAnchors and IntegrateData functions from Seurat. This integrated dataset was used for principal component analysis, variable gene identification, Shared Nearest Neighbor (SNN) clustering analysis, and Uniform Manifold Approximation and Projection (UMAP). Cell cycle phase genes are used to calculate cell cycle phase score for each cell which was used to regress out the effects of cell cycle heterogeneity.

Using the Louvain modularity optimization in Seurat, we explored clusters (unsupervised) at several resolution values, ranging from 0 to 1. Resolution is a parameter within the Louvain algorithm that controls the number of generated clusters. Marker genes for each cluster were identified by comparing gene expression from the target cluster against all other cells in the remaining clusters using the Wilcoxon rank sum test. Marker genes met the following criteria: 1) Log2 fold-change (Log2FC) ≥ 0.25 in the target cluster, 2) minimum of 25% of cells expressing the gene in both compared groups of cells, and 3) Bonferroni-corrected p-value < 0.05 [not applied]. A resolution value of 0.4 was ultimately selected, as distinct gene expression profiles were present in each cluster, while higher resolutions produced adjacent clusters with a high degree of overlapping marker genes.

Differentially expressed genes are identified using the Wilcoxon rank sum test with the following inclusion criteria: 1) minimum enrichment Log2FC ≥ 0.50 on either side; [not applied] 2) minimum percentage of cells expressing the gene ≥ 25 on either side; and 3) adjusted p-value < 0.05 [not applied].All plots were generated using various libraries in R v3.5.0-4.0.1(66), including Seurat, ggplot2 v3.2.1(67).

