## Supplemental figures S1-S4 for "Subset-specific and temporal control of effector and memory CD4+ T cell survival"

### Supplemental figures and legends

**A**

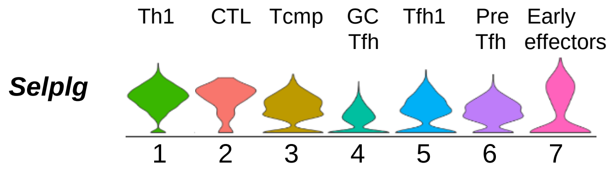


**B**

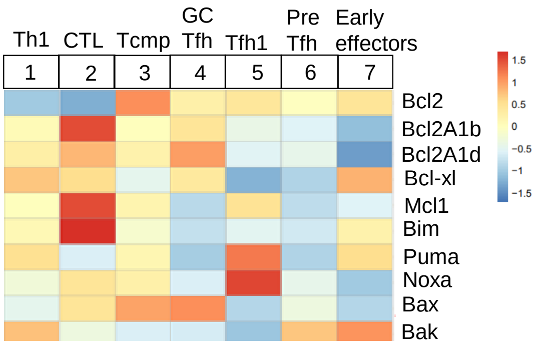


| Bcl2 family | Th1 | CTL | Tcmp | GC Tfh | Tfh1 | Pre-Tfh | Early effectors |
|---|---|---|---|---|---|---|---|
| Anti-apoptotics | A1b/d<br>BclxL<br>Mcl1 | A1b/d<br>BclxL<br>Mcl1 | Bcl2<br>A1b/d<br>Mcl1 | Bcl2<br>A1b/d<br>BclxL | Bcl2<br>Mcl1 | Bcl2 | Bcl2<br>BclxL |
| Pro-apoptotics | Bim<br>Puma<br>Bak | Bim<br>Noxa<br>Bax | Puma<br>Noxa<br>Bax<br>Bim | Bax | Puma<br>Noxa | Bak | Bim<br>Puma<br>Bak |

### Supplemental Fig 1

**scRNAseq of GP66-specific CD4+ T cells at d7 p.i. with LCMV** (A) Violin plot of *Selplg*(PSGL-1) expression across clusters (B) Heatmap of Z-scores for the average expression of Bcl-2 family members across clusters and table depicting the highest expressed anti and pro-apoptotic Bcl-2 family members across the different clusters.

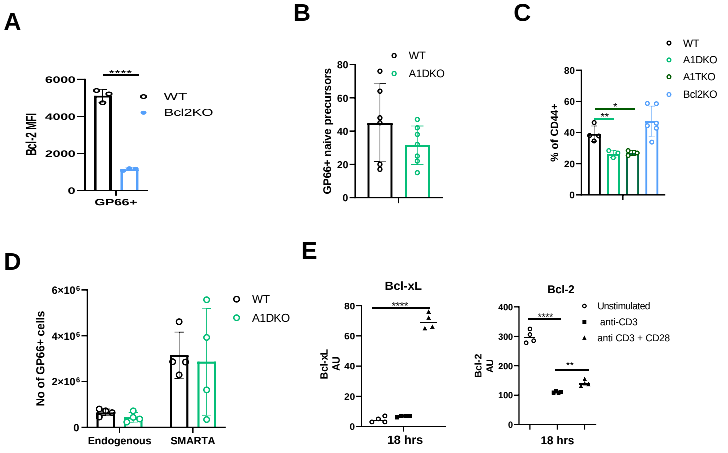


### Supplemental fig 2

(A) Efficient deletion of Bcl-2 in GP66-specific CD4+ T cells at d10 p.i. with LCMV (B) Naïve($CD44^{lo}$) GP6-specific CD4+ T cell precursors in the spleen of uninfected WT (open circle) and A1DKO (light green) mice (C) Frequency of CD44+ cells at d10 p.i. in WT (open circle), A1DKO (light green), A1TKO (dark green) and Bcl2KO (blue) mice(D) Splenic SMARTA CD4+ T cells were purified from congenic (CD45.1+) SMARTA transgenic mice by magnetic isolation.10,000 SMARTA CD4+ T cells were injected intravenously into WT and A1DKO mice, the mice were infected with LCMV the next day and sacrificed at d10 p.i. Number of splenic SMARTA cells in WT (open circle) and A1DKO (light green) recipients at d10 p.i. (E) Bcl-xL and Bcl-2 expression by qPCR in CD4+ naïve T cells magnetically enriched from spleens of WT mice and cultured without stimulation(open circles) or stimulated with anti-CD3 (1 μg/mL plate-bound) alone (squares) or in combination with anti-CD28(1 μg/ml)antibodies( triangles) for 18 hrs. Results are representative of at least 2 independent experiments with n=3 or more mice per group and show mean ± SD. *$p < 0.05$, **$p < 0.01$, Student's t test.

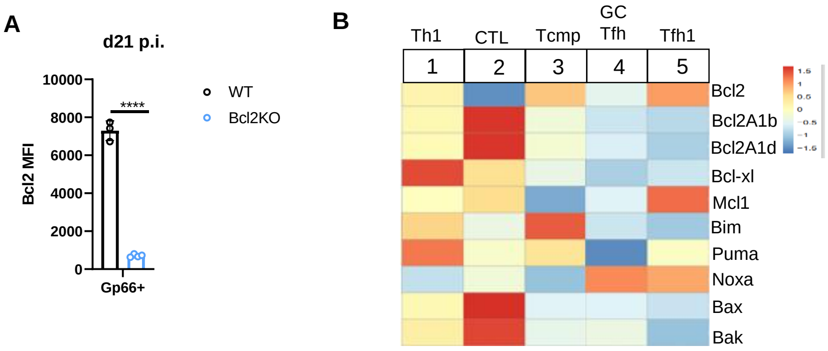


**Supplemental fig 3**

(A) Bcl2 expression in WT (open circles) vs Bcl2KO (blue) mice treated with tamoxifen from d11 to d15 post infection and sacrificed at d21 p.i. (B) Heatmap of Z-scores for the average expression of Bcl-2 family members across clusters from scRNAseq of GP66-specific CD4+ T cells at d24 p.i with LCMV. Results are representative of at least 2 independent experiments with n=3 or more mice per group and show mean ± SD. **** $p < 0.0001$ Student's t test.

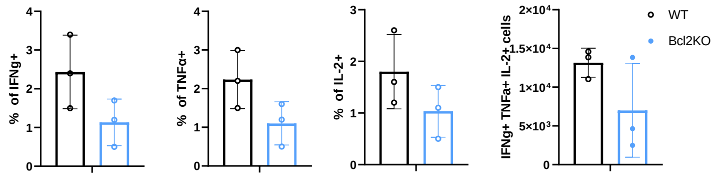


**Supplemental fig 4**

Bar graphs show the frequency of IL-2, TNFα and IFNγ positive cells and the cell numbers of triple producers of IL-2, TNFα and IFNγ in WT (open circle)and Bcl2KO (blue) at d90 p.i when Bcl-2 is deleted by tamoxifen administration at d85 to d89 p.i . Results are representative of at least 2 independent experiments with n=3 or more mice per group and show mean ± SD.
